# Postnatal increase in MRTF-dependent transcription reduces EGFR activation and proliferation of apical but not basal NSCs

**DOI:** 10.64898/2025.12.12.693930

**Authors:** Yujun Shen, Katja Baur, Martin Irmler, Johannes Beckers, Claudia Mandl, Gabriele Hölzl-Wenig, Francesca Ciccolini

**Author notes:** Correspondence should be addressed to: Dr Francesca Ciccolini, Department of Neurobiology, Interdisciplinary Center for Neurosciences (IZN), University of Heidelberg, Im Neuenheimer Feld 366, 69120 Heidelberg, Germany.

## Abstract

In the ventricular-subventricular zone (V-SVZ), both apical and basal NSCs undergo quiescence and proliferation. The epidermal growth factor receptor (EGFR) and interactions with the extracellular matrix are key regulators of adult NSC proliferation and quiescence, respectively. Here, we show that activation of EGFR significantly declines after the first postnatal weeks in apical NSCs. This decline is accompanied by a shift in serum responsive factor (SRF)-dependent transcription in apical NSCs. Specifically, growth-promoting genes targeted by SRF and ternary complex transcription factors are downregulated, whereas those targeted by SRF and myocardin-related transcription factors (MRTFs), including those involved in the extracellular matrix remodeling, are upregulated. Blocking of MRTFs, whose activity is regulated by the Rho/actin pathway and extracellular matrix interactions, restores EGFR activation and EGFR-dependent proliferation in adult apical NSCs. Thus, transcriptional programs regulated by extracellular cues differentially control EGFR activation and proliferation in neonatal and adult apical NSCs.

**Graphical abstract:** Graphical representation of the postnatal changes in transcriptional programs observed in apical and basal NSCs in V-SVZ.

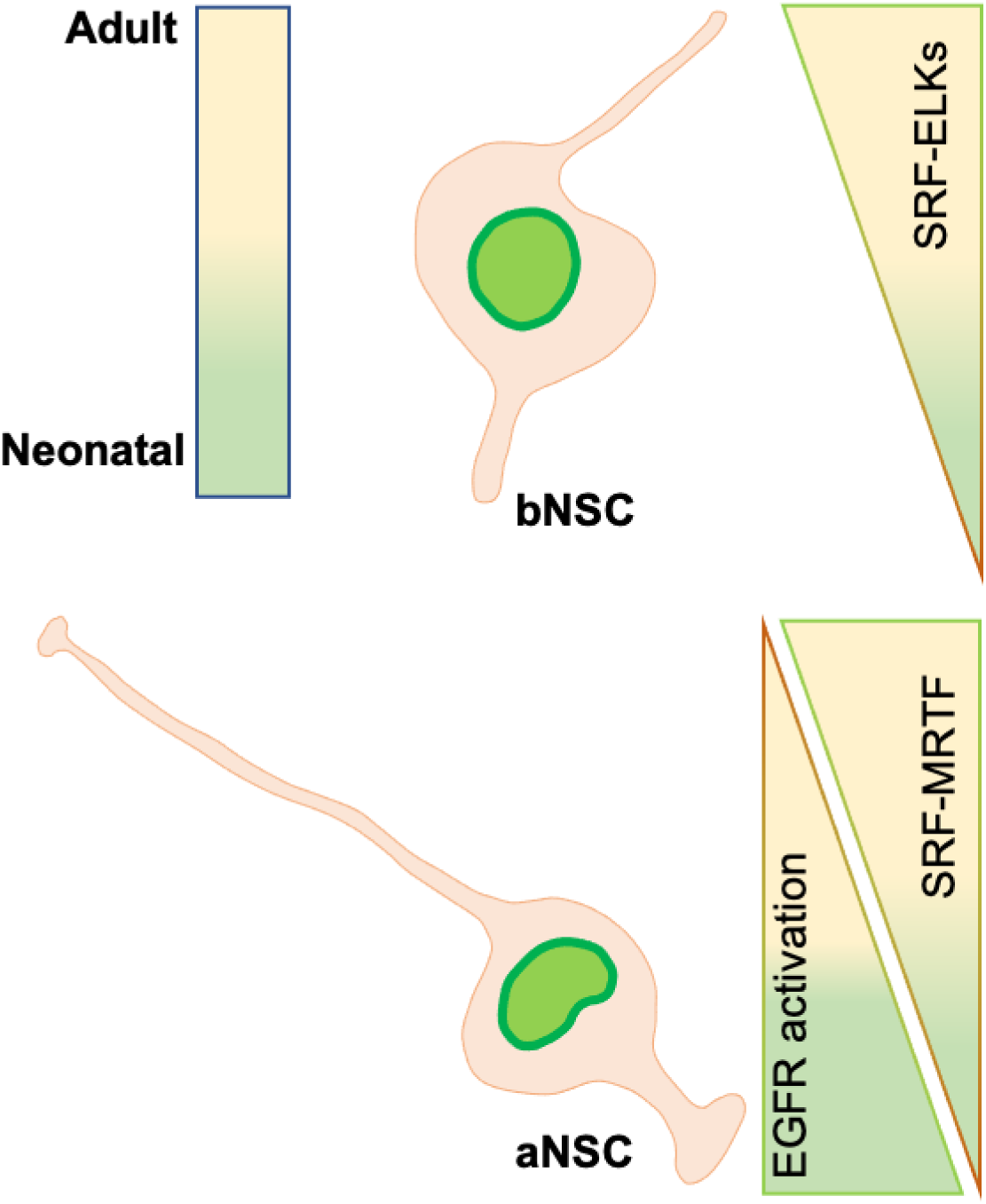

## Introduction

The generation of new olfactory bulb interneurons persists throughout adulthood and is sustained by neural stem cells (NSCs) residing in the ventricular/subventricular zone (V-SVZ) ^1^. We have recently demonstrated that adult NSCs in the V-SVZ comprise two distinct populations, i.e. apical and basal NSCs, which reside in different compartments of the niche and exhibit unique morphological and functional features ^2^. Apical NSCs, unlike the basal counterpart, display a radial glia-like cytoarchitecture with an apical membrane contacting the lateral ventricle, they largely exhibit Prominin immunoreactivity (P^+^) and frequently extend a primary cilium. In normal conditions, apical NSCs very rarely undergo division and do not significantly contribute to neurogenesis. Moreover, for at least the first six months after birth they represent the less abundant type of NSCs. Although the total number of both apical and basal NSCs declines with age, the strongest decrease is observed within the first two postnatal weeks especially in apical NSCs.

From mid-development onward, EGF acts as a potent NSC mitogen and, at later developmental stages, activation of EGFR is strongly associated with proliferation also *in vivo* ^3^ and it is required to sustain the proliferation of apical neural progenitors ^2, 4–7^. However, the mitogenic effect of EGF on apical P^+^ NSCs declines rapidly after birth, and in the adult V-SVZ is limited to a population of cells expressing high levels of EGFR ^2, 6, 8^. Underlying its centrality in NSC proliferation, EGFR interacts with several environmental signals like notch, beta integrin, and various neurotransmitters to regulate proliferation of adult NSCs in a manner that varies according age-dependent cues ^9–11^. In the case of the γ-aminobutyric acid (GABA), it promotes the proliferation of neonatal NSCs by altering EGFR trafficking downstream of GABA_A_Rs activation ^12–14^. However, the activation of GABA_A_Rs appears to have an opposite effect in the adult V-SVZ ^15^.

It is long known that expression levels as well as duration and intensity of EGFR activation can be associated with differential activation of MAPK, which in turn is important for the regulation of NSC proliferation ^16^. However, the molecular mechanisms underlying the different functions of EGFR according to age and cell type-dependent cues are still poorly understood. In this context, also the effect of transcriptional changes is scarcely investigated. For example, although it is known that quiescence induces expression of genes involved in the interaction with the extracellular matrix (EM)^17^, whether transcription affects also EGFR signalling is not known. In this regard, the serum response factor (SRF) is particularly interesting, because it can be activated via the Ras/MAPK or the Rho/actin cascade, thereby leading to different transcriptional outcomes. Mechanistically, this change in transcription involves the generation of alternative transcriptional complexes either with those belonging to the ternary complex factor (TCFs) or with the myocardin-related transcription factors (MTRFs) ^18^.

In this study we show that apical and basal NSCs display differences not only in the expression of genes associated with EGFR activation and signal transduction, but also in genes associated with SRF-dependent transcriptional response. We also show that many of these differences arise during the first two postnatal weeks and that they underlie the decreased ability of older apical NSCs to undergo EGFR-dependent proliferation compared to the neonatal counterpart.

## Matherials and Methods

### Animals and tissue dissection

Experiments with animals were carried out upon the receipt of permission from the local authorities and were done following the guidelines of laboratory animal use and care from Regierungspräsidium Karlsruhe and Heidelberg University. Neonatal mice were sacrificed by decapitation. Diazepam was administered by intraperitoneal injection as described before ^12–14^. Depending on the experiments, older mice (wild-type, C57Bl/6, and hGFAPtTA;H2B-GFP) were sacrificed by neck dislocation following CO_2_ inhalation or subjected to transcranial perfusion after being anesthetized ^13^. Briefly, mice were given two intraperitoneal injections (i.p. every 12 hours with 3 µg/g of body weight diazepam or PBS as control. When indicated, animals received an additional single intraperitoneal injection of 100 mg/kg of body weight of the thymidine analogue iododeoxyuridine (IdU) at the same time of the first diazepam/PBS injections. Mice were sacrificed 24 hours or 7 days following the first diazepam injection. For hGFAPtTA;H2B-GFP, 50 mg/ml of doxycycline was administered to 1 month old mice in the drinking water for the period of 4 weeks ^2^.

### Drugs

Diazepam (3µg/gr body weight). Mice were given two i.p. injections within 24 hours. PBS was used to similarly inject control mice. Muscimol (50 µM; Sigma). PD158780 (20 µM; Calbiochem/Merck, #513035). CCG-1423 (300 nM; Enzo, #BML-EI394-0010).

### Cell culture and flow cytometry

Subventricular zones from 8-week-old mice were dissected in dissection solution containing 150 mM sucrose, 125 mM NaCl, 3.5 mM KCl, 1.2 mM NaH_2_PO_4_, 2.4 mM CaCl_2_, 1.3 mM MgCl_2_, 6.65 mM glucose and 2 mM HEPES. The dissected tissue was digested with papain solution (1.6mg/ml) for 3 min at 37°C. Afterwards, the enzyme was inhibited with ice cold ovomucoid solution (1.4mg/ml) and washed away, and the tissue was dissociated in sort medium containing 50% NS-A medium, 50% Leibowitz L15 Medium, 2% B27 supplement, 1% FCS,1.33% D-(+)-Glucose, 10 ng/ml huFGF-2, 0.001% DNase, 2mM L-Glutamine and 100μg/ml penicillin/streptomycin. The cell suspension was then filtered in a 35 μm nylon mesh BD falcon tube to obtain a single cell solution. The cell suspension was incubated for 30 minutes at 4°C with an antibody against Prominin1 directly conjugated to an allophycocyanin (APC) fluorophore (Supplementary table S1) and/or EGF coupled to Alexa 647. After the incubation time, the non-bound antibody was washed away from the cell suspension by centrifugation for 5 minutes at 1000 rcf. Thereafter, cells were analysed and/or sorted using a FACSAria III flow cytometer (Becton and Dickinson). Sorting gates were set as previously illustrated ^19^.

### RNA isolation and quantitative PCR

Cell populations were sorted by FACS directly into RNA lysis buffer, vortexed for homogenization and stored at −80°C for RNA extraction. PicoPure™ RNA Isolation Kit (Applied Biosystems) was used to isolate RNA according to the manufacturer’s protocol. RNA concentration was measured using Denovix spectrophotometer. Reverse transcription and quantitative real-time PCR were done as previously explained ^19^. RNA-Seq was outsourced to BGI. The following Taqman probes were used: Egfr Assay ID Mm00433023_m1, Lrig1 Assay ID Mm00456116_m1, Aqp4 Assay ID Mm00802131_m1.

### RNA-seq analysis

RNA-Seq was done at BGI using the DNBSEQ Eukaryotic Strand-specific mRNA library protocol to generate PE100 data. Raw reads were filtered by the BGI standard pipeline to obtain cleaned data. Data filtering included removing adaptor sequences, contamination and low-quality reads from raw reads. Reads were aligned to the genome (Ensembl release 109 annotation on GRCm39) using STAR (v2.7.10b) and gene-level read counting was done with summarizeOverlaps (mode = ‘IntersectionNotEmpty’, package GenomicAlignments v1.28.0). Statistics was done with DESeq2 v1.40.2 including multiple testing by the Benjamini-Hochberg method. Classification of genes for Fig. 1 was based on the evaluation of the current literature and the depicted fold-changes were calculated by DESeq2 from the analysis of either neonatal or adult samples or from the combined analysis. Samples were processed separately for neonatal and adult samples and RNA-seq data has been submitted to the GEO database at NCBI (GSE311415).

**Figure 1.**
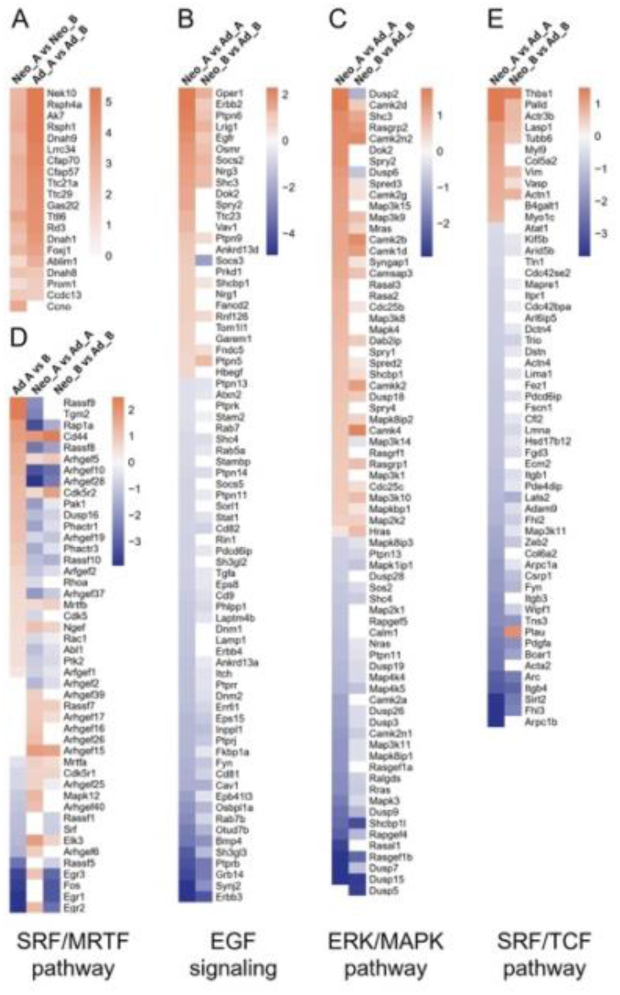
Changes in gene expression in neonatal and adult apical and basal NSCs. Heatmaps of the log2 fold-changes (log2FC) of genes in apical (A) NSCs and basal (B) NSCs isolated from the ventricular-subventricular zone (V-SVZ) at postnatal 1 week (Neo) and 3 weeks (Ad). Genes were significantly (padj<0.1) regulated in at least one comparison per heatmap (A) Cilia-associated genes including Prominin-1 in apical and basal NSCs at each age. (B) Genes associated with EGF signaling, including EGF superfamily receptors (EGFR, ERBB2, ERBB3, ERBB4) and key negative regulators of EGFR phosphorylation (e.g., ERFFI1, PHLPP1, EPB41L3). (C) Genes associated with ERK/MAPK pathway (e.g., DUSP3, DUSP6, DUSP7, RASAL1). (D) Genes associated with the SRF/TCF transcriptional pathway (e.g., SRF, ELK1, ELK4, MRTFA, MRTFB), MRTFs activation (e.g., TGM2) and downstream immediate early genes (e.g., FOS, EGR2, EGR3). (E) Genes targeted by the SRF/MRTF transcriptional pathway, particularly those involved in cytoskeleton organization and extracellular matrix interaction (e.g., ITGB1).

### Vibratome sectioning

Brains were embedded in 4% low melting agarose in a squared plastic scaffold. Once the agarose was solid, brains were glued to the slicing base of a Vibratome HM 650V. Coronal sections of 300 µm were made from the olfactory bulbs until the hippocampus. Slices were collected and kept in slicing sucrose contain μg/ml bicuculline at 4°C. Then, the slices were immediately transferred to a 24-well plate containing insert (Millicell-CM) incubated with muscimol or CCG-1423. 24 hours later, they were processed for Immunofluorescence staining.

### Immunocytofluorescence

After sorting, cells were plated on chamber slides (Nunc) coated with growth factor-reduced matrigel (Becton Dickinson) in Neurobasal-A medium (GIBCO) supplemented with B27, 1% fetal calf serum, in the presence of 2 ng/ml FGF-2. Cells were left to adhere to the substrate for one hour in the incubator before being fixed and processed for immunocytochemistry as described before ^6^. Primary antibody binding was detected using Alexa Cy3, Alexa 555-Alexa 647 conjugated secondary antibodies (Molecular Probes).

### Immunohistofluorescence

Postnatal day 7 (P7) mice were anaesthetized and perfused with a solution of 4% paraformaldehyde (PFA). Brains were removed and left in 4% PFA at 4°C. After additional 24 hours in 20% sucrose, brains were washed in PBS and embedded in 3% agarose in PBS. Coronal vibrotome sections were cut at a thickness of 70-100 μm and then processed for immunochemistry. Staining was revealed using Alexa 488 and PE-conjugated secondary antibodies, as previously described ^2^.

### Statistical analysis

For analysis of immunohistochemistry images of the dorsal, lateral and ventral regions of the V-SVZ were taken from each slice. For each animal a total of 3 slices were analyzed. Immunopositive cells were counted and normalized by total number of cells with DAPI stain or by region of interest (ROI) of a fixed area (15000 µm^2^). Images were acquired using a Leica SP8 confocal microscope using a HyDTM detector. Fiji/ImageJ software was used for measuring fluorescence intensities and normalization to background.

Data analysis was carried out using GraphPad Prism 5.0. Student’s t-test (two-tailed) or Analysis of Variance (ANOVA) were used with Welch’s correction when necessary (when variances showed significant difference according to the F-test).

## Results

### Changes in gene expression in apical and basal NSCs

We recently reported that during the first six months after birth, the most prominent decrease in the number of NSCs occurs within first two postnatal weeks, particularly among apical NSCs ^2^. We also showed that early after birth Prominin1 immunopositive (P^+^) NSCs largely lose their ability of to undergo EGF-elicited clone formation and that genes associated with EGF signalling are upregulated in basal compared to apical adult NSCs ^2^. To begin to investigate the mechanisms underlying the change in NSC number and its relationship with EGFR responsiveness, we performed RNA sequencing from apical and basal NSCs isolated from 1- and 3-week-old hGFAPtTA;H2B-GFP mice. Apical and basal NSCs were identified by nuclear tagging with the fusion protein H2B-GFP driven by a minimal human glial fibrillary acidic protein (hGFAP) promoter and were distinguished based on differential Prominin1 expression. Using a stringent cutoff of twofold change, we confirmed the selectivity of our isolation procedure as at both ages Prominin-1 and other genes associated with primary cilia were more expressed in apical than in basal NSCs (Fig.1A; all data shown in Fig. 1 is available in supplementary table S2). Moreover, upon comparisons of younger and older NSCs there were more cilia-related genes up-than down-regulated both in apical (32 vs 25) and in basal (20 vs 14) NSCs (Supplementary table S2B), which is in line with our previous findings that the number of primary cilia decreases in time. Next, we analyzed changes in the expression of genes associated with the EGF superfamily and EGFR intracellular transduction (Fig.1B). Using a cutoff of 1.5, we found that several genes associated with the negative regulation of EGFR phosphorylation at multiple levels were upregulated in older apical NSCs (Fig. 1B). This analysis revealed that the expression of all four receptors for the EGF superfamily was regulated within the first three postnatal weeks especially within the group of apical NSCs. For example, age led to a downregulation of EGFR and ERBB2 in both NSC groups, the effect was greater in apical than in basal NSCs (Supplementary table S2B). Age also led to an upregulation of the other receptors of the superfamily, i.e. the kinase dead ERBB3 and ERBB4, leading to increased expression in older NSCs and this effect was greater in apical than in basal NSCs (Fig1 B). A similar regulation was observed for several phosphatases like Ptprr, Ptprb, Ptprk, Ptpn13, and Phlpp1 ^20^ and for Sh3gl3, Sh3gl2, Eps15, Dnm1, Dnm2 and Rin1 coding for adaptor proteins involved in EGFR internalization and trafficking like ^21, 22, 23^ (Fig. 1B and Supplementary table S2B). Further highlighting changes in EGFR trafficking, compared to the neonatal counterpart and adult basal NSCs, the expression of genes involved in its internalization and sorting to lysosomes like Ankrd13a ^24^, Synj2 ^25^, Rab7, Laptm4b Rab7b, Lamp1 and Osbpl1a ^26–29^ was higher in three-week-old apical NSCs than in the younger group (Fig. 1B and Supplementary table S2B). Besides receptor trafficking, among the upregulated genes were the tumor suppressor protein Epb41l3 ^30^, preventing EGFR activation, and Socs5, Cd81, CD82 and CD9, which all provide a negative feedback inhibition of EGFR^31–33^ and Fkbp1a, associated both with the inhibition of EGFR phosphorylation ^34^ and receptor internalization^35^ (Fig. 1B and Supplementary table S2B). Instead, BMP4 which suppresses EGFR signalling^36^ and negatively regulate apical neural progenitor proliferation ^11^ and the leucine-rich repeats and immunoglobulin-like domains 1 (Lrig1), which is considered a NSC marker and a main regulator of EGFR activation in adult NSCs ^37^, were regulated during aging by more than two-fold both in apical and in basal NSCs leading to similar expression levels in the two adult NSC groups (Fig. 1B and Supplementary table S2B). Verification of gene expression analysis were performed by quantitative RT-PCR and immunohistochemistry (Supplementary Fig. S1A-C). Supporting the hypothesis that age leads to a decreased activation of EGFR in apical NSCs, immunohistochemistry with an antibody recognizing EGFR phosphorylated on tyrosine 1068 (pEGFR) revealed fewer pEGFR^+^ cells in the older V-SVZ especially at the apical side of the niche (Supplementary Fig. S1B). Activation of EGFR can be transduced through multiple pathways and the ERK/MAPK pathway by EGFR is essential to promote proliferation ^16^. Besides direct inhibitors of EGFR activation, using the same criteria we found also several modulators of the ERK/MAPK cascade differentially regulated in apical and basal NSCs. For example, the expression of several dual-specificity phosphatases (DUSP) ^20^ and the GTPase activating RASAL1 ^38^ was increased, whereas several genes of Sprouty/Spred family ^39^ were downregulated (Fig.1 C and Supplementary table S2C) during aging in apical but not basal NSCs. This indicates that transduction downstream of the EGFR differs between apical and basal NSCs. Activation of different transduction intracellular pathways downstream of EGFR can elicit distinct transcriptional responses. The activation of the ERK/MAPK cascade promotes the interaction of ternary complex factors (TCFs) the ELK1, 3, 4 and the serum responsive factor (SRF), thereby leading to the expression of c-fos and several early growth response genes (Egr) mediating cell cycle progression ^40, 41^. In contrast, changes in the actin cytoskeleton downstream of the Rho family of small GTPases promote association of SRF and the myocardin related transcription factor (MRTF) A and B. We found that Srf was less expressed in adult apical NSCs than in the basal counterpart but also that ElK3 was significantly downregulated during aging in apical NSCs, whereas Fos and several Egrs were upregulated in adult basal NSCs and downregulated in their apical counterpart (Fig.1 Dand Supplementary table S2D). In addition, some genes promoting MRTFs activation were upregulated in apical NSCs. These included transglutaminase 2 (TGM2) ^42^ and CD44 ^44^ which regulate NSC proliferation ^43^ and quiescence ^45^, respectively. Finally, several genes regulated by SRF/MRTF transcription were upregulated during aging in apical but not in basal NSCs (Fig.1 E and Supplementary table S2E). These included genes involved in the regulation of the cytoskeleton and the interaction with the extracellular matrix like Integrin beta 1 (ITGB1) ^43, 46–48^. Consistent with our expression analysis, immunostaining confirmed that ITGB1 expression levels increased at the apical side of the V-SVZ and in the adult brain (Supplementary Fig. S1B, C). Taken together, these data indicate that EGFR expression activation and downstream signaling are differentially regulated in apical and basal NSCs during the first three weeks after birth. In particular, the changes observed during the first weeks after birth appear associated in apical NSCs with a downregulation of EGFR activation and downstream ELK-mediated transcription.

### Diazepam administration leads to proliferation of basal but not apical NSCs in the adult V-SVZ

Next, we asked whether age-dependent changes in the activation and transduction of EGFR also are associated with a change in its ability to promote proliferation in apical NSCs. To test this possibility, we investigated whether GABA_A_R activation promotes apical NSC proliferation in the adult V-SVZ as in the neonatal counterpart ^12, 14^. We chose this experimental paradigm as we previously demonstrated that GABA_A_R activation promotes proliferation of neonatal P^+^ apical NSCs by a mechanism involving AQP4-mediated osmotic swelling, which leads to increased EGFR expression ^12, 14^. Since the RNAseq analysis revealed no major age-dependent changes in the expression of GABA_A_Rs and AQP4 in the two adult NSC populations (Supplementary table S2G and Fig. S1A), we hypothesized that apical and basal NSCs should be similarly able to respond to GABA_A_R activation. Therefore, we gave 8W old mice a first intraperitoneal injection of diazepam and IdU followed by a second intraperitoneal injection of diazepam alone. After further 24 hours or 7 days, we sacrificed the animals for further analysis (Supplementary Fig. S2A). Immunostaining of coronal brain sections from mice sacrificed at the earliest time point showed increased Ki67 and IdU immunoreactivity in diazepam-treated animals compared to controls (Supplementary Fig. S2B, C, left panel). However, the percentage of double positive cells was not affected, indicating that the treatment does not affect cell cycle dynamics but only cell cycle entry (Supplementary Fig. S2C, right panel). In mice sacrificed at 7days, only the number of total Ki67^+^ cells was affected by the treatment (Supplementary Fig. S2D, left panel). Furthermore, at this time point a greater portion of IdU^+^ cells were represented by IdU^+^/Ki67^+^ double positive cells (Supplementary Fig. S2 S2D, right panel), indicating that they were slow-dividing neural precursors.

We next used hGFAP;GFP-H2B mice to directly investigate the effect of diazepam injection on adult apical and basal NSC proliferation. For this analysis the two NSC groups were distinguished based on the distance of their fluorescent nuclei from the lateral ventricle as previously reported ^2^. Animals were administered doxycycline in the drinking water for one month to increase the percentage of quiescent NSCs within the group of basal NSCs ^2^ or left untreated (Fig. 2). This analysis revealed that both in untreated (Fig. 2A, B, left panel) and in doxycycline-treated animals (Fig. 2C, D, left panel) diazepam significantly increased the number of cycling H2B-GFP tagged (G^+^ Ki67^+^) NSCs. However, separate analysis of apical and basal NSCs showed that the increase in proliferation was significant only in basal but not in apical NSCs (Fig. 2 B, D, right panels). Notably, the number of mitotic basal NSCs, identified by the expression of phospho-Histone H3 (pHH3), revealed only a non-significant increase after the diazepam treatment (% basal G^+^ pHH3^+^ doxycycline-untreated: PBS = 0,75 ± 0.42, diazepam = 1.63 ± 0.91; doxycycline-treated: PBS = 0,61 ± 0.61, diazepam = 3.07 ± 1.88; n ≥ 3), which is consistent with our analysis above, indicating that diazepam promotes the proliferation of slow dividing NSCs. Taken together, these data show that in the adult V-SVZ, diazepam injection promotes the proliferation of slow dividing basal but not apical NSCs.

**Figure 2.**
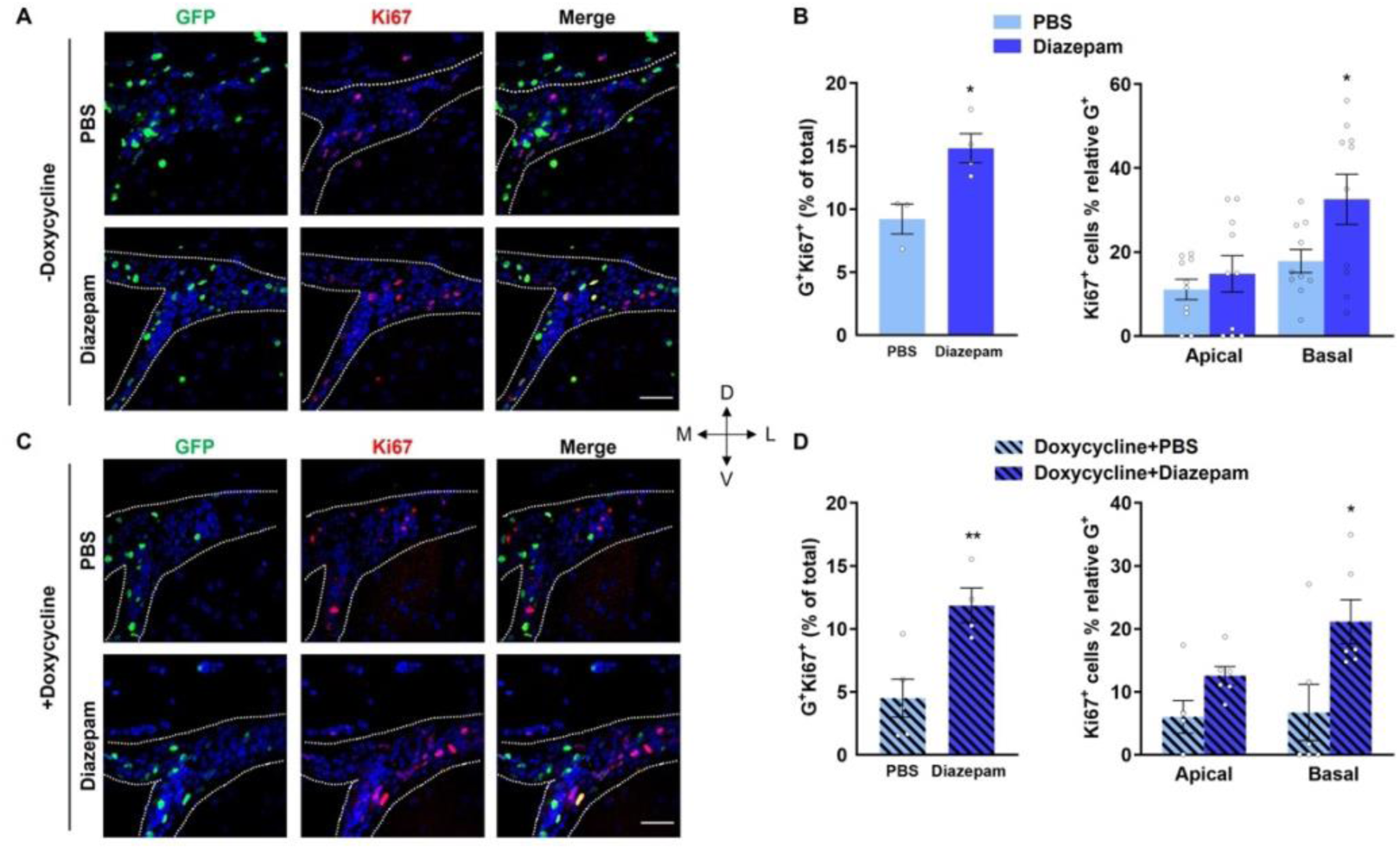
Diazepam promotes proliferation of basal but not apical NSCs. Adult mice were untreated (A, B) or given doxycycline in the drinking water (C, D) for a month. Thereafter, they were injected intraperitoneally with diazepam or PBS and perfused 24 hours later. (A, C) Confocal microphotographs of coronal sections from the SVZ of hGFAP-H2BGFP mice treated with doxycycline as indicated. Scale bar indicates 50 μm. (B, D) Quantification of total (left panel) and apical and basal cells double positive for H2BGFP (G^+^) and Ki67 (right panel). Apical and basal NSCs were identified based on their position respect to the ventricle cavity. Bars represent mean ± SEM. *Indicates significance: *P < 0.05, **P < 0.01, determined by unpaired Student’s t-test.

### Involvement of EGFR activation in the GABAergic regulation of adult NSC proliferation

Next, we investigated whether EGFR is required to induce proliferation upon GABA_A_R activation also in adult NSCs, as was previously observed in the neonatal V-SVZ ^12^. Firstly, we measured the effect of the GABA_A_R agonist muscimol on the cell size (Supplementary Fig. S3A, B) and the expression level of EGFR at the membrane (Supplementary Fig. S3C, D) of isolated apical and basal NSCs. Consistent with our RNAseq expression analysis, which revealed similar expression of GABA_A_Rs in the two cell groups, the cell size both of NSC populations was increased upon muscimol treatment (Supplementary Fig. S3A, B). However, the treatment led to an increase in EGFR expression only in adult basal NSCs, but not in the apical counterpart (Supplementary Fig. S3C, D), which is in contrast with our previous observation in neonatal P^+^ NSCs ^12, 14^. Besides EGFR expression, changes in cell morphology may affect interaction with the surrounding matrix, which is important for the regulation of quiescence ^49^. Since ITGB1 is important for maintaining quiescence of adult NSCs and may regulate EGFR activation ^9^, we next investigated the effect of diazepam on EGFR phosphorylation and ITGB1 expression, in the two NSC groups by immunohistochemistry (Fig. 3A). Independent of doxycycline administration, quantitative analysis revealed lower levels of receptor phosphorylation in adult apical NSCs than in the basal counterpart (Fig. 3B), consistent with our hypothesis that age leads to an impairment of EGFR activation in apical NSCs. Furthermore, treatment with diazepam led to a significant increase in pEGFR immunoreactivity in basal but not in apical NSCs (Fig. 3B, upper panel). Diazepam treatment led also to a decrease in Itgb1 transcript (Fig. 3B, middle panel) and protein (Fig. 3B, lower panel) expression in adult apical NSCs. However, this effect was only observed in the V-SVZ of mice had been treated with doxycycline. Notably, a significant increase in pEGFR was observed only in basal G^+^/ITGB1^-^ cells upon diazepam injection (Supplementary Fig. S4A, B). Similar changes in ITGB1 immunoreactivity were also observed upon examination of the fraction of cycling NSCs (Supplementary Fig. S4C, D). Taken together, these results indicate that increased EGFR activation is necessary for diazepam to affect basal NSCs proliferation, whereas changes in ITGB1 expression alone are not enough to promote NSC proliferation upon GABA_A_R activation. Consistent with this conclusion, pharmacological blockade of EGFR with PD158780 (PD) in basal NSCs prevented the increase in proliferation elicited by exposure to muscimol (Fig. 4A, B).

**Figure 3.**
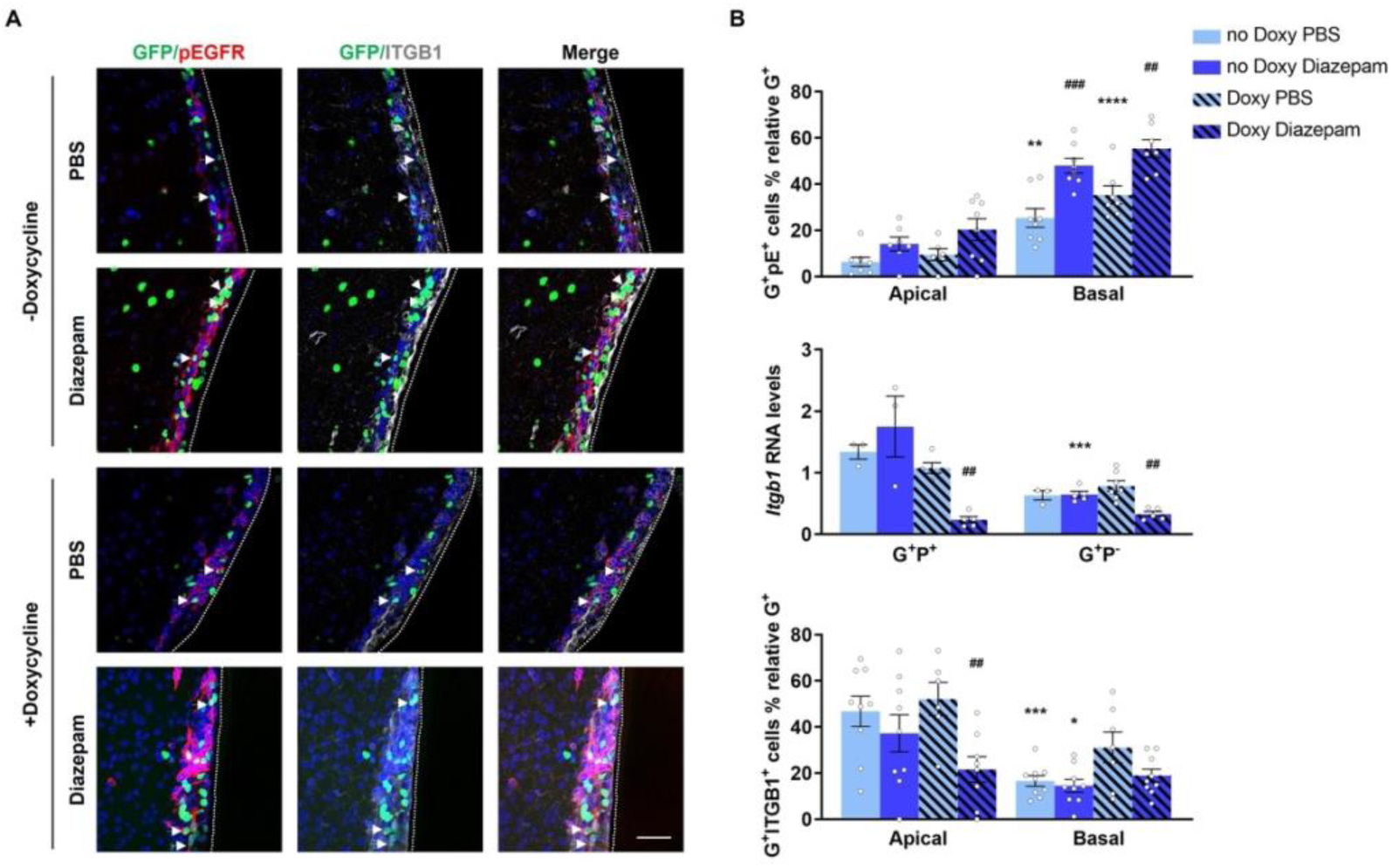
Changes in EGFR phosphorylation but not in ITGB1 expression are associated with the proliferation inducing effect of Diazepam on NSCS. Measurement of phosphorylated EGFR and ITGB1 protein and transcript expression on brain slices and in sorted apical (G^+^P^+^) and basal (G^+^P^-^) NSCs. Mice were sacrificed and analysed 24h after being intraperitoneally injected with diazepam or PBS as control. (A) Representative confocal microphotographs of the V-SVZ stained of adult GFAP-H2B mice with antibodies for phosphorylated EGFR (pEGFR) and ITGB1, treated with doxycycline as indicated. Scale bar indicates 50 μm. (B) Quantitative analyses of the number of total apical and basal G^+^ cells displaying pEGFR immunoreactivity (upper panel), relative Itgb1 mRNA levels (middle panel) and quantitative analyses of apical and basal G^+^ cells dispaying ITGB1 immunoreactivity (bottom panel). Data shown as mean ± SEM, *= significant difference between corresponding apical (G^+^P^+^) and basal (G^+^P^-^); #= significant difference between control and treatment. *p<0.05, ***p<0.001, #p<0.05, ##p<0.01, ####p<0.0001.

**Figure 4.**
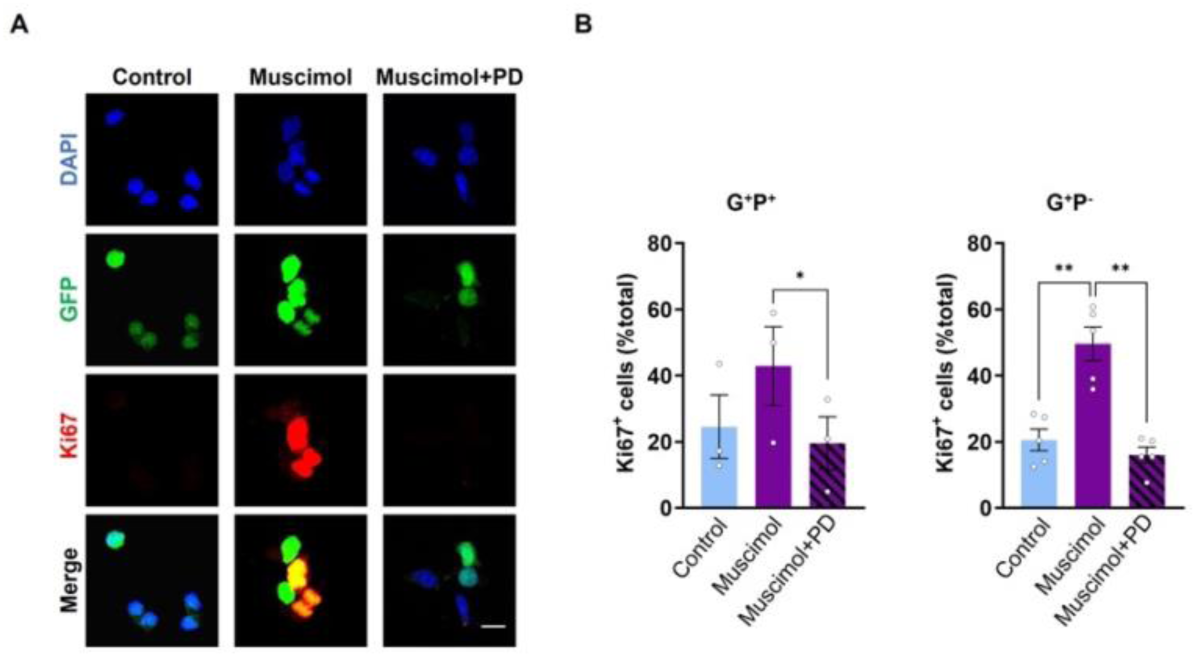
Blockade of EGFR activation prevents the increase in basal NSC proliferation elicited by exposure to muscimol. Analysis of Ki67 expression in isolated apical (G^+^P^+^) and basal (G^+^P^-^) NSCs treated for 24 hours with muscimol in the presence or absence of PD158780 (PD) as indicated. Scale bar indicates 10 μm. (A) Representative confocal microphotographs of G^+^P^-^ basal NSCs upon staining as indicated. (B) Quantitative analyses of the number of Ki67 immunopositive cells in apical (G^+^P^+^) and basal (G^+^P^-^) NSCs cultures treated as indicated. N≥3. Bars represent mean ± SEM. *Indicates significance: *P < 0.05, **P < 0.01 determined by one-way ANOVA with Tukey’s multiple comparisons test.

Thus, our data show that both apical and basal NSC undergo swelling in response to GABA_A_R activation. However, unlike what it was previously reported in neonatal NSCs, activation of the neurotransmitter receptor elicits EGFR overexpression and phosphorylation only in basal but not in apical NSCs. This underscores the decreased ability of adult apical NSCs to transduce EGFR signalling compared to their neonatal counterpart.

### GABA_A_R activation promotes proliferation of apical NSCs upon blockade of MRTF-dependent transcription

We next used the small molecule CCG-1423 to directly inhibit MRTF-mediated transcription ^50^ to investigate whether it contributes to restricting EGFR activation in adult NSCs. Acute brain slices were exposed to CCG-1423 and/or muscimol in the culture medium or left untreated as a control. After 24 hours the effect of the treatments on apical and basal NSC proliferation and EGFR phosphorylation was assessed by immunostaining (Fig. 5A). In contrast to treatment with muscimol alone, blocking MRTF-mediated transcription with the concomitant activation of GABA_A_Rs caused an increase in the number of apical NSCs (Fig. 5B, upper left panel) and a significant increase in Ki67 expression (Fig. 5B, upper right panel) in apical NSCs. Instead, none of the treatments significantly affected the number of basal G^+^ cells (Fig. 5E). Independent of the presence of CCG-1423, exposure to muscimol increased the number of cycling basal G^+^ cells, whereas treatment with CCG-1423 alone did not significantly affect proliferation (Fig. 5G).

**Figure 5.**
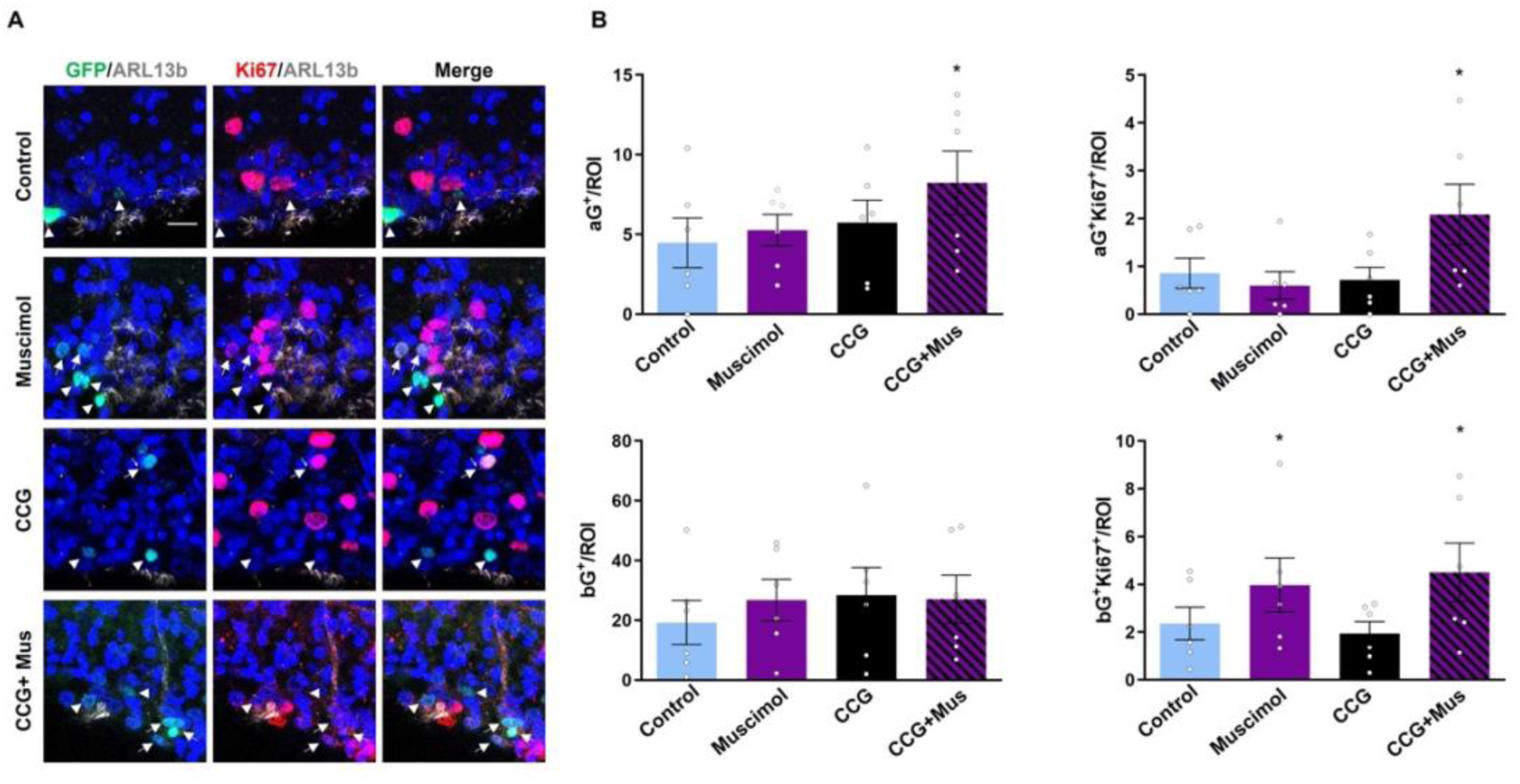
Activation of GABA_A_R and simultaneous MRTF blockade increased the proliferation of apical NSCs. Analysis of the effect of the Muscimol and/or CCG-1423 treatment of organotypic slices of the V/SVZ obtained from adult hGFAP-H2BGFP mice on the total number apical and basal NSCs and on Ki67 expression within the groups of G^-^ and G^+^ apical and basal cells. (A) Representative confocal microphotographs of coronal sections of the V-SVZ treated as indicated and immunostained for Ki67 (red) and ARL13b (gray). Scale bar indicates 10 μm. (B) Number of total G^+^ (upper left panel) and G^+^KI67^+^ (upper right panel) apical cells and total G^+^ (bottom left panel) and G^+^KI67^+^ (bottom right panel) basal cells per region of interest (ROI). Bars represent mean ± SEM. *Indicates significance: **P < 0.01, ****P < 0.0001 determined by one-way ANOVA with Tukey’s multiple comparisons test.

Importantly, increased cycling of G^+^ cells was coupled to increased EGFR phosphorylation (Fig. 6) This was observed both in apical (Fig. 6A, B upper right panel) and basal (Fig. 6B lower left panel) cells in slices treated with both CCG-1423 and muscimol, and only at the basal side of the niche in slices treated with muscimol alone (Fig. 6B lower right panel). Again, blockade of MRTF alone did not significantly alter the phosphorylation of the receptor.

**Figure 6.**
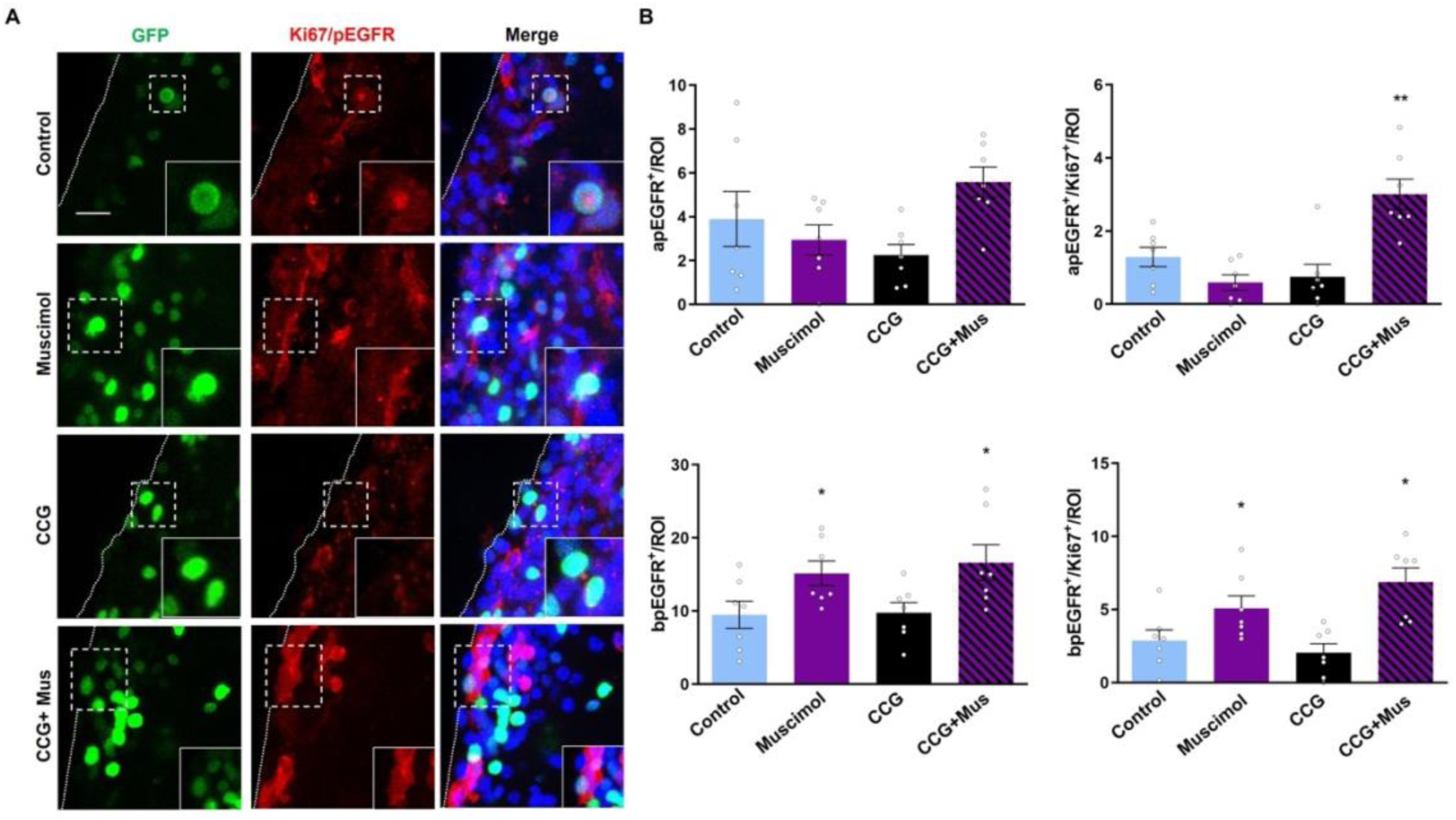
Activation of GABA_A_R and simultaneous MRTF blockade increased EGFR phoshorylation in apical NSCs. Analysis of the effect of the Muscimol and/or CCG-1423 treatment of organotypic slices of the V/SVZ obtained from adult hGFAP-H2BGFP mice on EGFR phosphorylation in non-cycling and cycling apical and basal NSCs (A) Representative confocal microphotographs of coronal sections of the V-SVZ treated as indicated and immunostained for Ki67 (nuclear red) and pEGFR (cytoplasmic red). Scale bar indicates 10 μm. (B) Number of apical (upper panels) and basal (lower panels) G^+^ NSCs per region of interest (ROI) displaying both nuclear and cytoplasmic (Ki67^+^/pEGFR^+^; right panels) or only cytoplasmic (Ki67^-^/pEGFR^+^; left panels) fluorescent signal. Bars represent mean ± SEM. *Indicates significance: *P < 0.05, **P < 0.01, ***P < 0.001, ****P < 0.0001 determined by one-way ANOVA with Tukey’s multiple comparisons test.

Taken together, our data show that the postnatal increase in MRTF-mediated transcription in apical NSCs counteract the ability of EGFR to activate and promote apical NSC proliferation.

## Discussion

We report here for the first time that SRF/MRTF transcriptional activity increases with postnatal age in apical but not in basal NSCs, thereby affecting the ability of EGFR to activate and promote proliferation of adult apical NSCs. A diagram illustrating the changes underlying the differential regulation of EGFR activation in neonatal apical and basal NSCs is illustrated in figure 7.

**Figure7.**
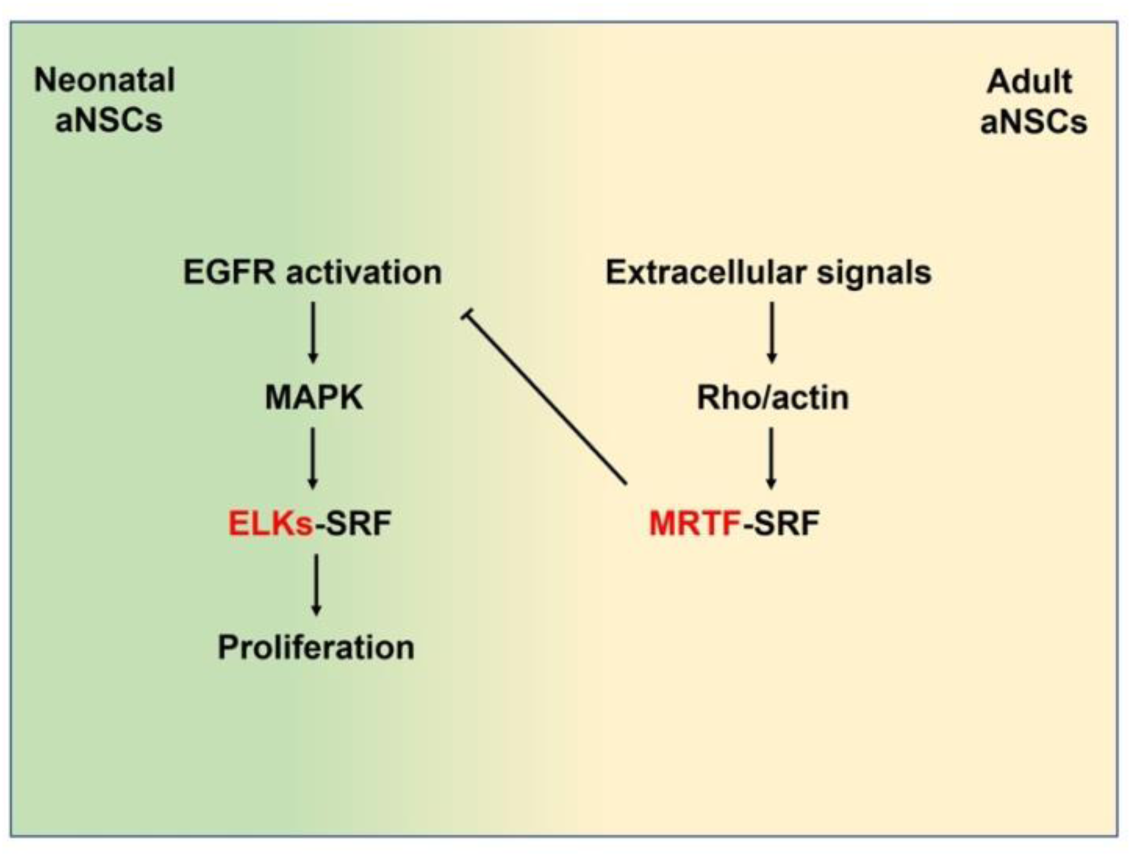
Diagram illustrating postnatal changes in SRF dependent transcription in apical NSCs and their effect on EGFR activation and EGFR-dependent proliferation.

Two types of astrocytes have been identified in the postnatal V-SVZ so called Type B1 and type B2 cells ^51^. Type B 1 astrocytes have characteristics of apical NSCs like for example apical radial glia morphology. They also have been considered the only population of NSCs in the adult V-SVZ. Type B2 cells were also originally defined essentially based on morphological characteristics established by electron microscopic examination and have been mainly considered niche astrocytes ^51^. However, since a subset of type B2 cell was proliferating, it was possible that they included also more undifferentiated progenitors. Indeed, we recently could show that a subset of these basal cells not only have intrinsic characteristics of NSCs, but they also represent the greatest NSC population and the major source of olfactory bulb interneurons ^2^. These findings were recently confirmed by a study showing that type B2 cells, devoid of an apical membrane contacting the lateral ventricle, represent the main population of neurogenic NSCs in the adult V-SVZ ^52^. However, in other aspects this latter study reached conclusions contrasting with ours. For example, the frequency of primary cilia observed in basal NSCs and type B2 cells are very different. Whereas virtually every type B2 cells displays a primary cilium we reported that only 5% of basal NSCs does. Furthermore, type B1 and B2 cells were found to express similar levels of Prominin1, which is in contrast with our findings that less than 10% of basal NSCs display Prominin1 expression at the cell surface. Importantly, that extension of primary cilia is a hallmark difference between Prominin1 immunopositive and immunonegative cells has been already reported ^6, 53^. Moreover, by means of direct single cell observation, we could also show that the vast majority of primary cilia are present in Prominin1 immunopositive and not immunonegative cells ^6^. Taking all this into account, a likely explanation for the discrepancies between the two studies, is that Arantxa Cebrian-Silla et al. only focused on the analysis of the small subset of type B2 cells which correspond to the small fraction of basal NSCs expressing high levels of Prominin-1. This interpretation would be also consistent with the observation that differentially expressed genes between type B1 and B2 cells code mainly for protein associated with the apical and basal cytoarchitecture and no significant differences in Prominin expression ^51, 52^.

In contrast, in the present study we could highlight that the two group of cells not only are characterized by different expression levels of Prominin1 and genes coding for ciliary proteins, but they also differ in the expression of multiple genes affecting the EGFR transduction/transcription axis.

We have here tested how the changes in EGFR related gene expression affect EGFR transduction and NSCs proliferation in the context of GABA_A_R activation. Although, several studies have previously highlighted that GABAergic signaling modulates the proliferation of neural progenitors, the direction of the observed changes and the mechanisms associated are not consistent. At least in part, the inconsistency reflects differences in membrane properties between different neuronal progenitor types, which in turn affects the cellular response ^3, 54^. For example, multiple reports highlighted that GABA_A_R signalling promotes depolarization and calcium signalling in intermediate precursors leading to cell cycle exit in the embryonic SVZ ^55–58^. Endogenous GABA_A_R activation in the adult V-SVZ appeared to have a similar effect on transit amplifying progenitors ^59^. Previous studies on NSCs also come to different conclusions. For example, it was observed that GABA_A_R promotes proliferation of radial glia in the embryonic brain ^56, 58^ and in neonatal NSCs ^12, 60^. However, an opposite effect of GABA_A_R on proliferation was observed in embryonic stem cells ^61^ and in NSCs in the adult V-SVZ ^15^. Our current study provides an explanation for these controversial observations by showing that although GABA_A_R activation induce osmotic swelling in both apical and basal adult NSCs, this leads to increased proliferation only in the latter, where swelling is accompanied by recruitment of EGFR. These observations are consistent with our previous findings that EGFR activation underlies the GABAergic effect on NSC proliferation^12, 14^. They also highlight that neonatal and adult apical NSCs differ in their ability to respond to EGFR signalling.

Notably, our findings underscore that the changes in EGFR signalling involve the regulation of EGFR activation rather than solely a change in EGFR expression. As a matter of fact, several intrinsic and extrinsic factors other than EGFR expression levels can affect responsiveness in NSCs ^62, 63^. We here show that also MRTF-dependent transcription contributes to this regulation and that MRTF blockade restores the ability of GAB_A_AR to promote EGFR phosphorylation and proliferation of apical NSCs. In light of our finding that the SRF/MRTF transcriptional axis is also increased in older apical NSCs compared to the neonatal counterpart, we conclude that this is responsible for the change in EGFR signalling observed between the two group of cells. The increased activation of MRTF-dependent transcription in apical NSCs could be due to either the increased expression of genes responsible for the activation of the rho/actin pathway, genes affecting the interaction with the extracellular matrix as well as genes regulating the composition of the glycocalyx, which affect the movement of molecule across the cellular membrane ^64^ and are differentially modified in apical and basal NSCs with increasing age ^11^. Supporting these conclusions, we found that age not only increased in apical NSCs the expression of genes responsible for the activation of the Rho/actin pathway, but also of genes like TGM2 affecting the stiffness of the extracellular matrix ^45^.

Thus, niche interaction and MRTF signalling regulate EGFR-dependent proliferation postnatal apical but not basal NSCs.

## Acknowledgements

We thank the Neurobiology department for providing a supporting environment.

