## supplementary figures for "Postnatal increase in MRTF-dependent transcription reduces EGFR activation and proliferation of apical but not basal NSCs"

### Supplementary Figure and Legends

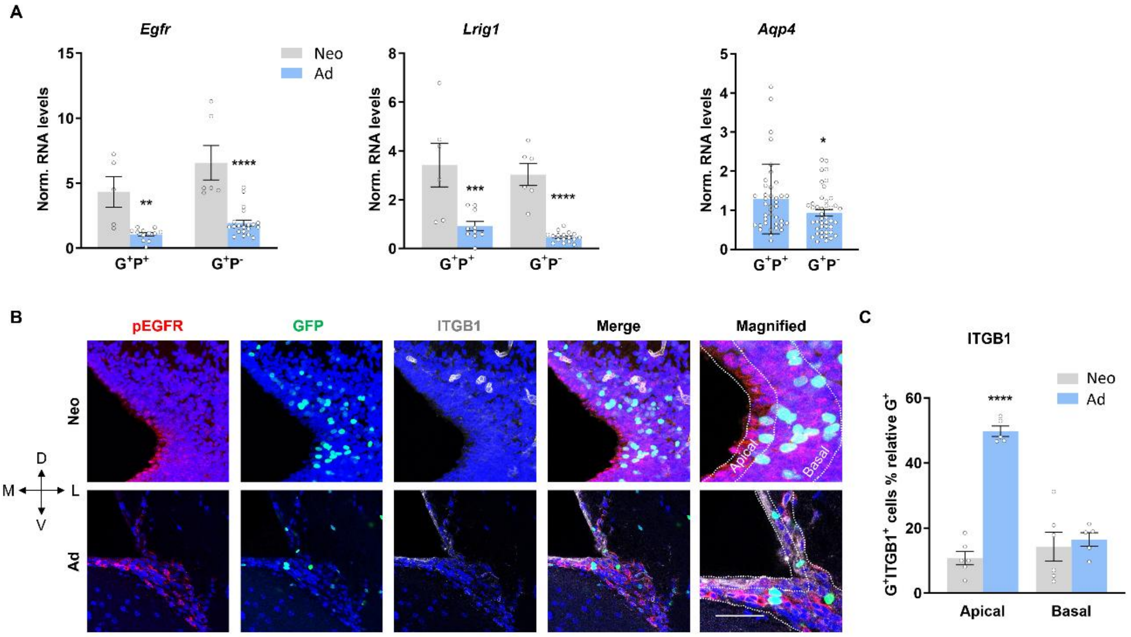


**Figure S1 Age-dependent changes in *Egfr* and *Lrig1* transcript levels, EGFR phosphorylation and ITGB1 expression in apical and basal NSCs.**

(A) Relative mRNA expression levels for the indicated genes.

(B) Representative confocal photomicrographs of coronal sections of the V-SVZ of neonatal (Neo; upper panel) or adult (Ad; lower panel) hGFAP-H2BGFP mice immunostained as indicated. Nuclear GFP expression and membrane pEGFR immunoreactivity are shown in green and red respectively. DAPI counterstaining of the nuclei is blue. Scale bar indicates 50 µm.

(C) Quantification of $GFP^+$ and/or $ITGB1^+$ cells in the V-SVZ, as a percentage of total DAPI cells N ≥ 5.

Bars represent mean ± SEM. *Indicates significance: *P < 0.05, **P < 0.01, ***P < 0.001, ****P < 0.0001 determined by one-way ANOVA with Tukey's multiple comparisons test (A, C) or two-tailed unpaired Student’s t-test.

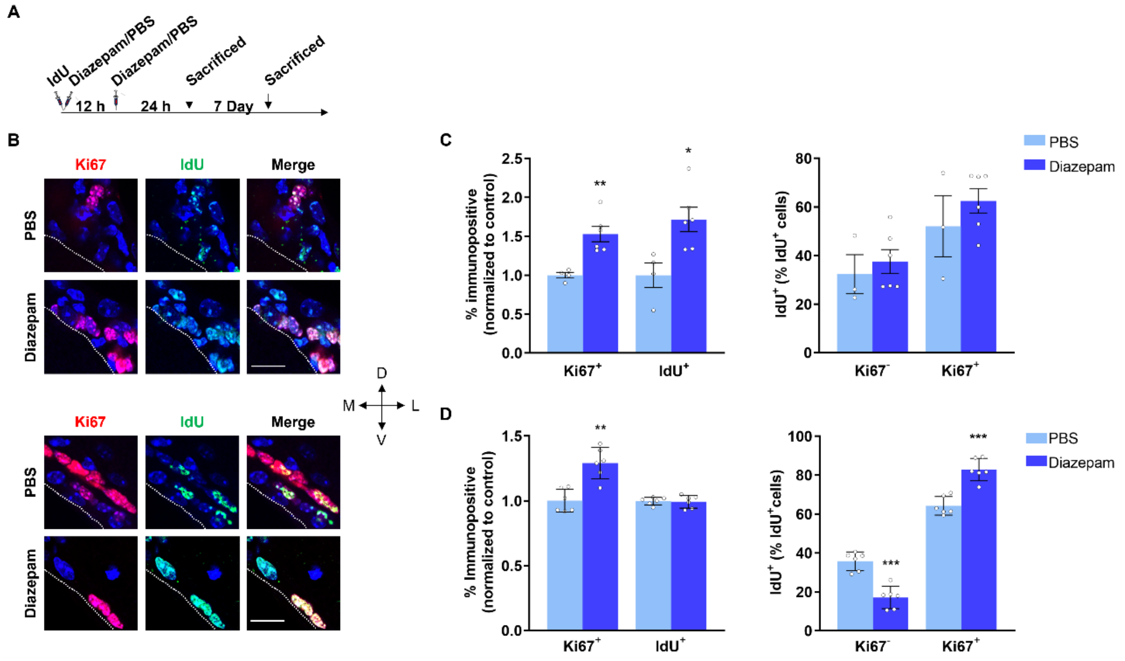


**Figure S2 Diazepam promotes proliferation in the adult V-SVZ.**

Analysis of Ki67 expression and IdU incorporation *in vivo*.

(A) Diagram illustrating the temporal sequence for IdU and diazepam administration by intraperitoneal injection.

(B) Representative confocal photomicrographs of coronal sections of the V-SVZ of mice sacrificed and immunostained for Ki67 and IdU 1 (upper panel) and 7 days (lower panel) after the first intraperitoneal injection. DAPI counterstaining of the nuclei is shown in blue. Scale bar indicates 20 μm.

(C-D) Quantitative analysis of the number of label-retaining cells in mice 1 (C, N ≥ 3) and 7 days (D, N= 6) after the first intraperitoneal injection.

Bars represent mean ± SEM. *Indicates significance: *P < 0.05, **P < 0.01, ***P < 0.001 determined by unpaired Student's t-test.

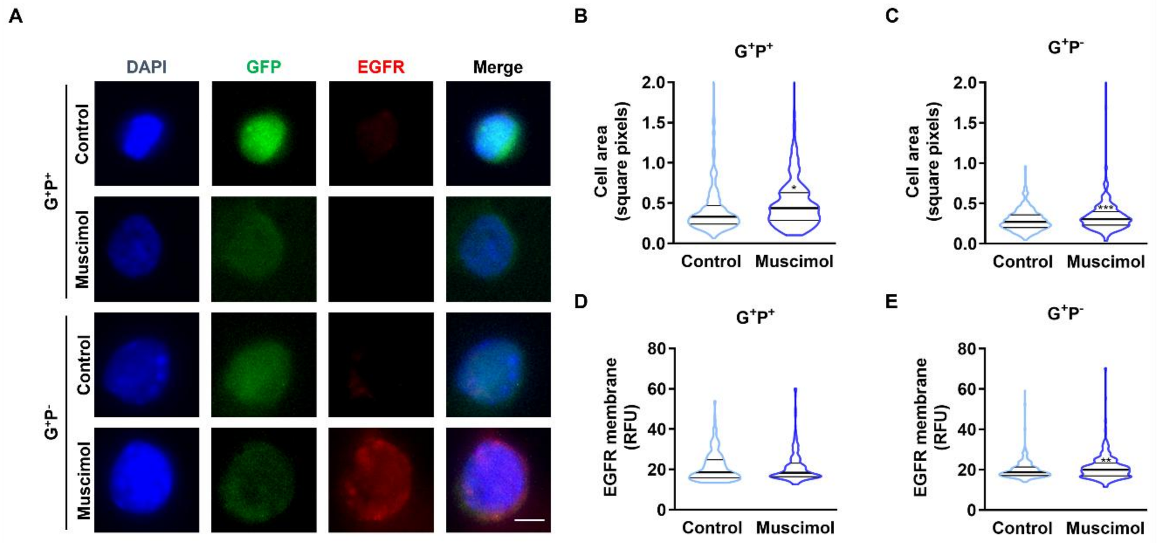


**Figure S3 GABA$_A$R activation promotes cells swelling of apical and basal NSCs and upregulation of EGFR expression only on the latter.**

Effect of muscimol on cell size and membrane expression of EGFR in isolated apical ($G^+P^+$) and basal ($G^+P^-$) NSCs.

(A) representative microphotographs of the given cell type treated as indicated after immunostaining for EGFR (red) and DAPI counterstaining of the nuclei (blue). Scale bar indicates 5 μm.

(B, C) The violin plots compare the distribution of cell area in square pixels for apical (B) and basal (C) NSCs cells exposed to muscimol or left untreated as control.

(D, E) The violin plots compare EGFR membrane expression in relative fluorescence units (RFU) for apical (D) and basal (E) NSCs cells exposed to muscimol or left untreated as control. N≥200 cells from N≥4 mice per condition (*$p<0.05$, **$p<0.01$, ***$p<0.001$).

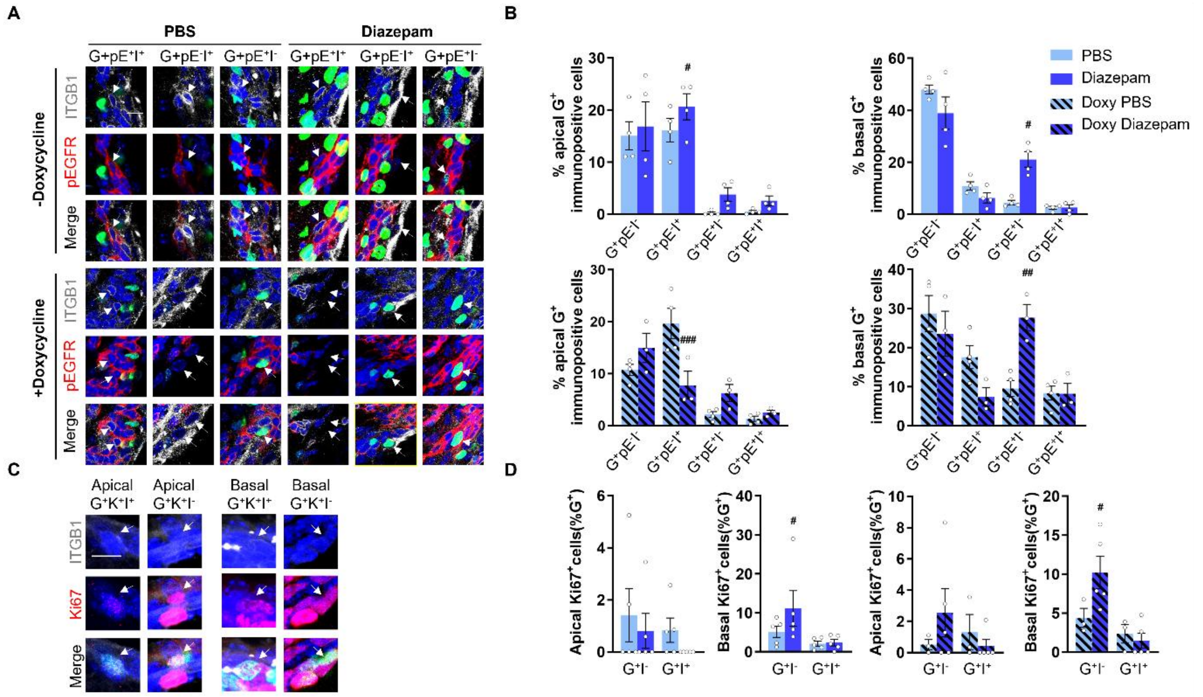


**Figure S4 Diazepam injection increases the percentage of basal but not apical NSCs displaying increased EGFR phosphorylation and lacking ITGB1 expression.**

Analysis of the changes in pEGFR (pE), ITGB1 (I) and Ki67 expression on coronal sections of the V-SVZ of adult hGFAP-H2BGFP mice sacrificed 24h after being intraperitoneally injected with diazepam/PBS. Before Diazepam administration some mice were treated with doxycycline for a month.

(A) Representative confocal microphotographs of coronal sections of the V-SVZ of adult hGFAP-H2BGFP mice, treated with doxycycline as indicated, upon double immunostaining for ITGB1 (gray) and pEGFR (red) and Dapi (blue) counterstaining of the nuclei. Scale bar indicates 10 µm.

(B) Quantitative analyses of the number of double positive cells for ITGB1 and pEGFR in apical (left panels) and basal (right panels) from untreated mice (upper panels) and mice that were given doxycycline (lower panels).

(C) Representative confocal microphotographs of coronal sections of the V-SVZ upon double immunostaining for ITGB1 (gray) and Ki67 (red) and Dapi (blue) counterstaining of the nuclei. Scale bar indicates 10 µm.

(D) Quantitative analyses of the number of double positive cells for ITGB1 and Ki67 in apical and basal NSC s as indicated.

Data shown as mean ± SEM, #p<0.05.
