## Supplementary table S1 for "Postnatal increase in MRTF-dependent transcription reduces EGFR activation and proliferation of apical but not basal NSCs"

**Suplementary Table S1**

Antibodies for flow cytometry

| Antibody | Fluorophore | Host | Dilution | Company, catalog # |
| --- | --- | --- | --- | --- |
| Prominin-1 | APC | Rat | 1:100 | Miltenyi Biotec, 130-102-197 |

Dying cells were reavealed by staining with propidium iodide (PI) (1μg/ml).

Antibodies for Immunohistochemistrry

| Antibody | Host | Dilution | Company, catalog # |
| --- | --- | --- | --- |
| Ki67 | Rabbit  Rabbit | 1:100  1:3000 | Abcam, 16667  Arigo,ARG11083 |
| BrdU (IdU staining) | Mouse | 1:10 | Hybridoma Bank, G3G4 |
| Phospho-EGFR (Y1068) | Rabbit | 1:500 | Abcam, 2220 |
| ITGB1 | Rat | 1:500 | Millipore, MAB1997 |
| ARL13b | Mouse | 1:500 | Biolegend, 857602 |
| PHH3 | Mouse |  | Invitrogen, MA5-15220 |

Staining with 4‘,6-Diamidine-2’phenylindole-dihydrochloride (DAPI) (0,1μg/ml) was used to counterstain the cell nuclei.
